# Mucosal Associated Invariant T (MAIT) Cell Responses Differ by Sex in COVID-19

**DOI:** 10.1101/2020.12.01.407148

**Authors:** Chen Yu, Sejiro Littleton, Nicholas Giroux, Rose Mathew, Shengli Ding, Joan Kalnitsky, Elizabeth W. Petzold, Hong Chung, Grecia Rivera Palomino, Tomer Rotstein, Rui Xi, Emily R. Ko, Ephraim L. Tsalik, Gregory D. Sempowski, Thomas N. Denny, Thomas W. Burke, Micah T. McClain, Christopher W. Woods, Xiling Shen, Daniel R. Saban

**Author notes:** These authors contributed equally to this work (Chen Yu, Sejiro Littleton).

## Abstract

Sexual dimorphisms in immune responses contribute to coronavirus disease 2019 (COVID-19) outcomes, yet the mechanisms governing this disparity remain incompletely understood. We carried out sex-balanced sampling of peripheral blood mononuclear cells from confirmed COVID-19 inpatients and outpatients, uninfected close contacts, and healthy controls for 36-color flow cytometry and single cell RNA-sequencing. Our results revealed a pronounced reduction of circulating mucosal associated invariant T (MAIT) cells in infected females. Integration of published COVID-19 airway tissue datasets implicate that this reduction represented a major wave of MAIT cell extravasation during early infection in females. Moreover, female MAIT cells possessed an immunologically active gene signature, whereas male counterparts were pro-apoptotic. Collectively, our findings uncover a female-specific protective MAIT profile, potentially shedding light on reduced COVID-19 susceptibility in females.

## MAIN TEXT

Severe Acute Respiratory Syndrome Coronavirus 2 (SARS-CoV-2) has led to a global pandemic of Coronavirus Disease 2019 (COVID-19) and a death toll of more than over 1.4 million people and rising (*1*). Among reported sex disaggregated data, males are disproportionately affected by SARS-CoV-2, with a higher incidence of cases, mortality, and morbidity (*2*). This follows a similar trend toward higher case fatality rates for males in Severe Acute Respiratory Syndrome Coronavirus (SARS-CoV) and Middle East Respiratory Syndrome Coronavirus (MERS-CoV), as well as with experiments using SARS-CoV mouse models (*2–6*). Sex differences in the immune response are thought to be a key contributing factor to these coronavirus disease outcomes, agreeing with the current body of knowledge that innate and adaptive immune responses are substantially altered across sex (*7–10*). Specific to SARS-CoV-2 infection, responses of both lymphocytes and myeloid cells were shown to be associated with COVID-19 outcomes (*11–19*). Correspondingly, a recent study on sex differences in COVID-19 immune responses uncovered an association between poor disease outcomes in males and weak T cell responses in both CD4^+^ and CD8^+^ compartments, whereas poor outcomes in females were associated with high innate immune cytokines, tumor necrosis factor superfamily (TNFSF)-10 and interleukin (IL)-15 (*20*). The sex differences elucidated in this seminal study further cement the need to better understand the mechanisms governing sex-specific susceptibility to SARS-CoV-2.

In the current study, we carried out sex-balanced sampling of peripheral blood mononuclear cells (PBMCs) from COVID-19 patients and control subjects for 36-color flow cytometry and single cell RNA-sequencing (scRNA-seq) analyses. A total of 88 samples were analyzed from 45 individuals. Details on subject demographics and sample information are summarized in (**Fig. 1A**, **table 1** and **S1**). Briefly, we analyzed samples from 28 patients with COVID-19 as confirmed by a positive SARS-CoV-2 PCR and/or IgG seroconversion. These included 9 inpatient subjects (20%), 7 requiring intensive care, henceforth referred to as “hospitalized.” An additional 19 subjects were identified in outpatient settings (42.2%), henceforth referred to as “infected”. Most of these COVID-19 confirmed cases were sampled longitudinally (a range 1-28 days) including pre- and post-anti-SARS-CoV-2 immunoglobin (IgG) seroconversion. The dates of symptom onset for all confirmed COVID-19 subjects were recorded at enrollment, providing an illness range of 1-40 days. We also recorded symptom severity, obtained via investigator survey on 39 symptoms related to COVID-19 (see Methods). Additionally, we included 7 subjects (15.6%) henceforth referred to as “exposed,” who were also sampled at multiple timepoints. These subjects, despite being close contacts of infected individuals, remained with negligible symptom scores, were negative for SARS-CoV-2 by PCR, and did not demonstrate detectable anti-SARS-CoV-2 IgG for at least 2 months after enrollment. Lastly, we included a group of 10 “healthy” subjects (22.2%) who were enrolled prior to the pandemic in 2019 and did not show any symptoms associated with COVID-19 or other respiratory illness (*21*).

**Fig. 1.**
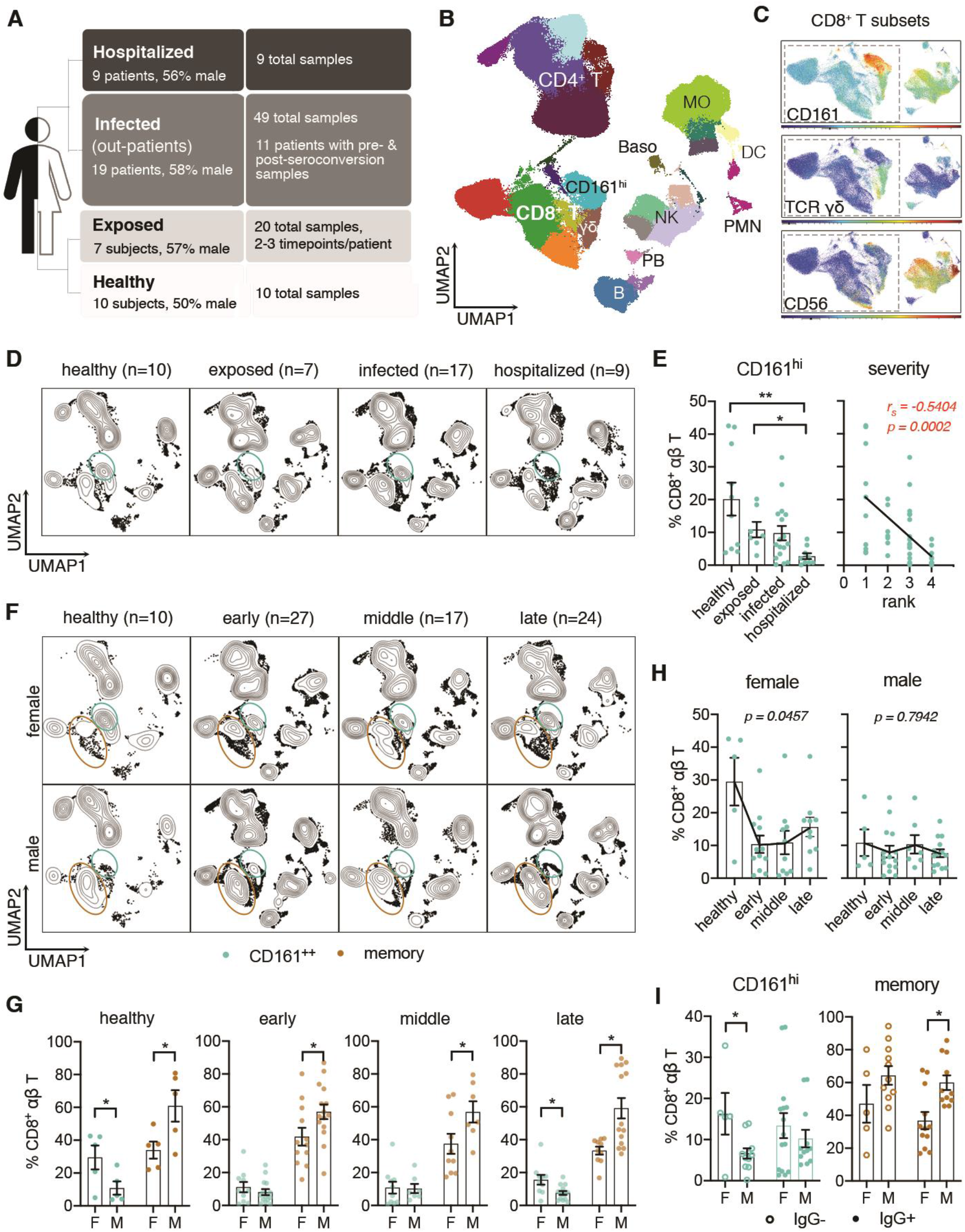
Sex-Specific CD8^+^ T cell Responses in PBMCs of COVID-19 Patients. (**A**) Overview of patient groups in this study. (**B**) UMAP visualization of PBMC subsets identified by FlowSOM clustering. Samples from all participants were pooled and down-sampled to 3,000 live CD45^+^ cells per sample. MO, monocytes; NK, natural killer cells; DC, dendritic cells, PMNs, polymorphonuclear neutrophils. Baso, basophils. (**C**) Expression of CD161, TCR γδ and CD56 in CD8^+^ T cell subsets. (**D**) UMAP of samples grouped by disease severity. Samples collected within 3 days from enrollment were included. (**E**) Frequencies of CD161^hi^ T cells in different severity groups (left) and their correlation with severity rank (right). (**F**) UMAP of samples stratified by sex and time post symptom onset (early, ≤14 days; middle, >15 days and ≤21 days; late, >22 days). (**G**) Frequencies of CD161^hi^ and memory CD8^+^ T cells between sex and timepoints. (**H**) Sex-specific changes of CD161^hi^ cells frequencies shown in G. (**I**) Frequencies of CD161^hi^ and memory CD8^+^ T cells in the samples from confirmed COVID-19 subjects pre- and post-seroconversion. Data were plotted as mean ± standard error. Significance was determined by Kruskal-Wallis test with Dunn’s test (E, H) or Mann Whitney test (G, I): \**p*<0.05, \*\**p*<0.01.

**Table 1.** Summary of Patient Demographics and Sample Information.

| Group |  | healthy | exposed | infected | hospitalized | total |
| --- | --- | --- | --- | --- | --- | --- |
| **Age**<br>mean ± SD (range) | | 39.70 ± 13.68<br>(25-61) | 42.57 ± 13.93<br>(17-60) | 36.73 ± 13.88<br>(20-65) | 59.44 ± 15.11<br>(31-76) | 42.84 ± 16.06<br>(17-76) |
| **# subjects**<br>(F:M ratio) |  | n = 10<br>(5:5) | n = 7<br>(3:4) | n = 19<br>(8:11) | n = 9<br>(4:5) | n = 45<br>(20:25) |
| **Race**<br>n (%) | African American | 3 (30.00%) | 0 | 1 (5.26%) | 5 (55.56%) | 9 (20.00%) |
|  | Asian | 0 | 1 (14.29%) | 2 (10.53%) | 0 | 3 (6.67%) |
|  | White | 7 (70.00%) | 5 (71.42%) | 16 (84.21%) | 3 (33.33%) | 31 (68.89%) |
|  | Others/<br>unknown | 0 | 1 (14.29%) | 0 | 1 (11.11%) | 2 (4.44%) |
| **Days** since onset<br>when enrolled<br>mean ± SD (range) | | N/A | 15.17 ± 11.25<br>(5-33) | 11.18 ± 4.30<br>(3-19) | 8.25 ± 6.14<br>(1-18) | 11.19 ± 6.73<br>(1-33) |
| **# samples** of flow<br>cytometry (F:M ratio) |  | n = 10<br>(5:5) | n = 20<br>(9:11) | n = 44<br>(21:23) | n = 9<br>(4:5) | n = 83<br>(39:44) |
| **# samples** of scRNA-seq (F:M ratio) |  | n = 5<br>(F:M = 3:2) | n = 8<br>(F:M = 3:5) | n = 29<br>(F:M = 12:17) | n = 6<br>(F:M = 1:5) | n = 48<br>(F:M = 19:29) |
| **Days** since onset<br>when samples collected<br>mean ± SD (range) | | N/A | 26.94 ± 14.94<br>(5-61) | 18.27 ± 8.44<br>(3-40) | 8.25 ± 6.14<br>(1-18) | 19.32 ± 11.44<br>(1-61) |

### Immune Profiling of COVID-19 Patient PBMCs Reveals Sex Differences in CD8^+^ Lymphocytes

With these flow cytometry data (**Table S2**), we generated a map of immune cell populations and their subsets by down-sampling to 3,000 viable CD45^+^ singlets per sample and concatenated all data for Uniform Manifold Approximation and Projection (UMAP) (*22*) and unbiased clustering via Flow Self-Organizing Maps (FlowSOM) (*23*). Unique marker expressions of respective populations facilitated our annotation of major PBMC populations, including CD4^+^ and CD8^+^ (αβ) T cells, γδ T cells, B cells, plasmablasts, natural killer (NK) cells, monocytes (MO), and dendritic cells (DC), and confirmed by manual analysis (**Fig. 1B**, and **fig. S1, A** to **E**). Sub-populations were also annotated in this manner, such as CD45RA^+^ CD27^+^ CCR7^+^ naive, CD45RA^−^ CCR7^+^ central memory (CM), CD45RA^−^ CCR7^−^ effector memory (EM) and CD45RA^+^ CD27^−^CCR7^−^ terminally differentiated effector memory (EMRA) CD8^+^ T cells, as well as CD8^+^ CD161^hi^ T cells and other indicated sub-populations (**Fig. 1B**, and **fig. S1, A** to **E**). We noted that a minor population of basophils (Baso) and neutrophils (PMN) primarily from hospitalized patients were detected (**Fig. 1B**), despite using a PBMC isolation protocol.

We next set out to examine our flow cytometry dataset for immune populations that exhibited major quantitative changes in COVID-19. Our first strategy was to stratify the data by disease severity (i.e., healthy, exposed, infected, and hospitalized). We noted that samples from hospitalized patients had substantially fewer PBMCs suggestive of lymphopenia (*24*). Manual gating of all flow cytometry events was performed for this analysis. Our results showed differences in B cells (naïve, IgD^+^ non-class switch, and plasmablasts); natural killer (NK) cells (CD56^lo^ populations); DCs (CD141^+^, CD1c^+^, and pDCs); monocytes (classical, intermediate and nonclassical); CD4^+^ (EM) and CD8^+^ (EM) αβ T cells (**Fig. S2**). Interestingly, our data also revealed a high statistical significance (*p*=0.0006) in CD8^+^ CD161^hi^ T cells (**Fig. S2**) prompting us to look closer at these cells. Regarding annotation of this CD161^hi^ cluster, because the overwhelming majority of events are low to negative for CD56 and for T-cell receptor (TCR)-γδ (**Fig. 1C**), the phenotype is largely consistent with mucosal associated invariant T (MAIT) cells, but not NKT or γδ T cells. This designation is congruent with recent work in COVID-19 PBMCs (*25–27*). We therefore conclude that the frequencies of certain myeloid and lymphocyte populations are affected in COVID-19, including a major effect on CD8^+^ CD161^hi^ T cells.

Knowing that the overwhelming majority of the CD8^+^ CD161^hi^ population (henceforth referred to as CD161^hi^) in our and others’ datasets (*25–27*) is likely comprised of MAIT cells, we performed a more focused analysis of this cluster in COVID-19 (**Fig. S3A**). First, we analyzed their frequencies by disease severity using the samples taken within 3 days of enrollment, the timepoint most proximal to the initial symptom score recordings. Results were displayed via UMAP contour plots, revealing a reduction in these cells in the SARS-CoV-2 settings (**Fig. 1D**). Manual gating from all flow cytometry events revealed a significant reduction when comparing healthy (p=0.0036) or exposed (p=0.0488) subjects versus hospitalized subjects, as well as a negative correlation (p=0.0002) with disease severity (**Fig. 1E** and **Fig. S3B**). We also characterized the frequencies of CD161^hi^ cells stratified by time post symptom onset, including early (≤ 14 days), middle (15 to 21 days), and late (>21 days) timepoints. In addition, we separated the data by sex given the known sex differences in immune responses in COVID-19 (*2, 20*). Results showed that within the CD8^+^ compartment of healthy subjects, females had greater frequencies of CD161^hi^ cells relative to males, whereas males had greater frequencies of CD8^+^ memory T cells (combined EMRA, EM and CM) (**Fig. 1F**, and **G**). No obvious changes of naïve CD8^+^ T were found (**Fig. S3C**). While the memory cell predominance in males was preserved at all timepoints, the greater abundance of CD161^hi^ cells in females was lost at early and middle timepoints (albeit not in late disease). The loss of this difference was due to a precipitous drop of CD161^hi^ cells in females at early and middle timepoints (**Fig. 1H**). Lastly, we stratified data from confirmed COVID-19 patients by seroconversion status. This showed CD161^hi^ cells were higher in females relative to males prior to seroconversion, whereas CD8^+^ memory cells were higher in males in seroconverted subjects (**Fig. 1I**). Taken together, we identified a female-specific decline in circulating CD161^hi^ cell frequencies upon exposure/infection of SARS-CoV-2. This sex-specific reduction may be due to extravasation into airway tissues, thereby suggesting a key sex-specific role for these CD161^hi^ cells in COVID-19.

### scRNA-seq of PBMCs Implicates Involvement of CD161^hi^ Lymphocyte Responses in COVID-19

Given the potentially important role for CD161^hi^ cells in COVID-19, we sought to further characterize this population by scRNA-seq (10x Genomics). We analyzed 48 different PBMC samples from 24 subjects across all groups (**Table 1** and **table S1**). Data were processed using Seurat 3 package (*28*) and subsequent transcript-based annotation was carried out (**Fig. 2A, fig. S4, A** and **B**). Focusing on the T cells in the data (**Fig. 2B** and **table S3**), we were able to identify CD161^hi^ cells in a single cluster containing high *KLRB1* (i.e., CD161) expression, and co-expression of *CD3D* and *CD8A*, as well as *TRAV1-2* (**Fig. 2C** and **fig. S4B**), which encodes the Vα7.2 invariant TCR alpha chain on MAIT cells. Grouping these data by disease severity showed that hospitalized patients had lower frequencies of T cells, including CD161^hi^ cells (**Fig. 2D**), agreeing with our flow cytometry findings and consistant with reported lymphopenia in severe COVID-19 patients (*11, 20, 29–32*). Also showing the same trend as our flow cytometry data was the high frequency of CD161^hi^ cells in healthy females (**Fig. 2E**), although it did not reach statistical significance due to the variations between healthy females and males. Next, to address the functional role of this CD161^hi^ cluster in COVID-19, we performed gene enrichment analysis using differentially expressed genes (DEGs). With several top ranked hits consisting of immune pathways and an estrogen-dependent pathway (**Fig. 2F**), our results inferred a sex-specific immune response of these CD161^hi^ cells in COVID-19. To further characterize functional inferences, we applied the CellphoneDB package (*33*) and analyzed ligand-receptor interactions with monocyte clusters within our data (**Fig. S5, A** and **B**), given the critical link that was previously published between monocyte activation in COVID-19 outcomes (*12, 13, 20, 34*). Our results inferred unique interactions between CD161^hi^ cells and monocytes with the following gene-pairs: *KLRB1_CLEC2D*, *CCL5_CCR1*, *CXCR6_CXCL16*, and *IL18_IL-18R* (**Fig. S5, C** and **D**). Moreover, the number of interaction-counts of monocytes was the most abundant with the CD161^hi^ cluster relative to all major T cell populations (**Fig. S5E**). Taken together, these transcriptome findings further support a significant role for circulating CD161^hi^ cells in the SARS-CoV-2 immune response.

**Fig. 2.**
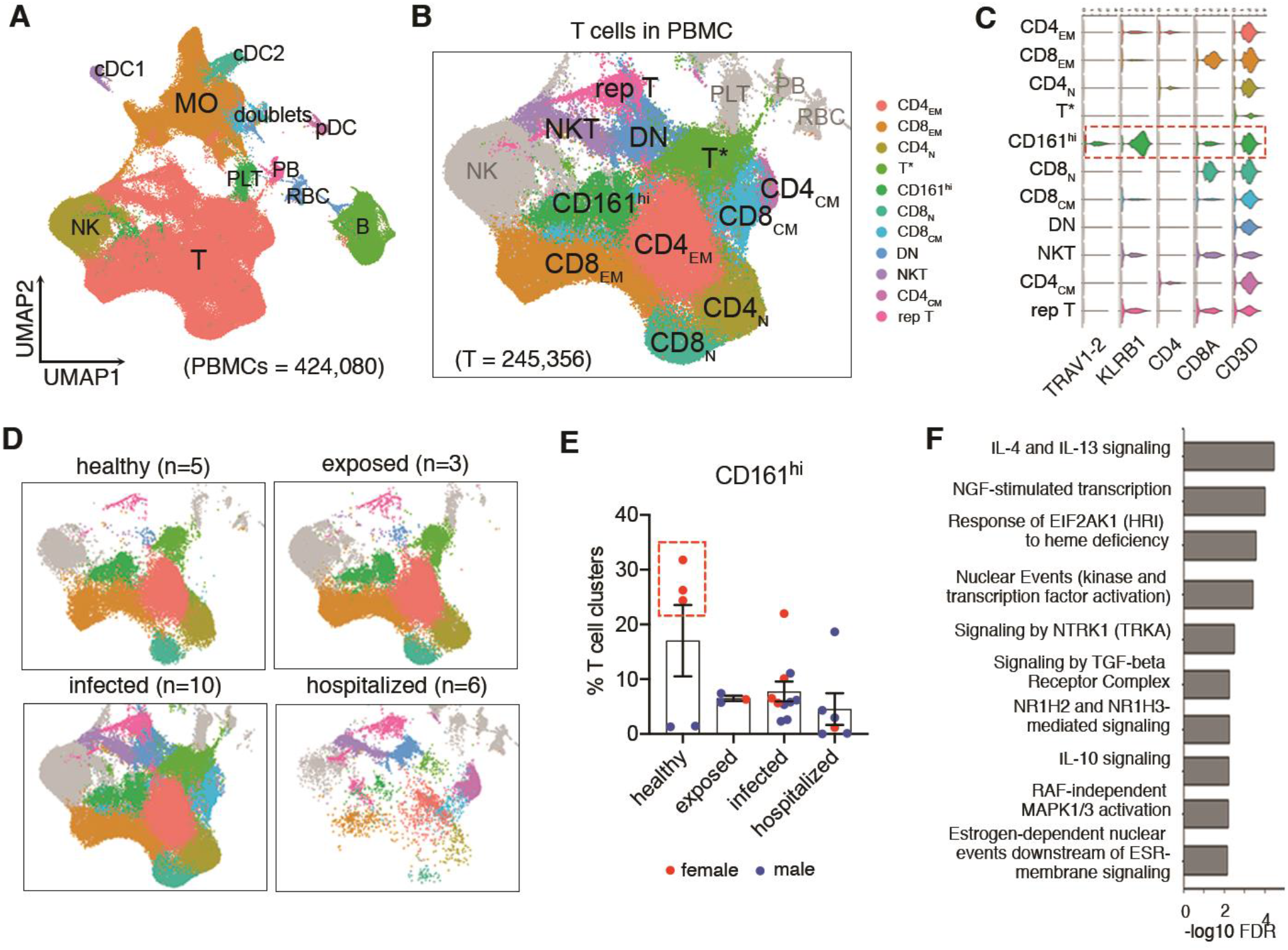
Characterization of CD8^+^CD161^hi^ T cells in COVID-19 using scRNA-seq. (**A**) UMAP and unsupervised cluster analysis of PBMCs. MO, monocytes; RBC, red blood cells; PB, plasmablasts; PLT, platelets; DC, dendritic cells. (**B** and **C**) Visualization of different T cell subsets with high resolution in UMAP (B) and expression of their marker genes as indicated in Violin plots (C). T* cluster likely represents a dropout population with low UMI counts. N, naïve; EM, effect memory; CM, central memory; DN, double negative; rep, replicating. (**D**) Changes of T cell subsets with disease severity. N, the number of individuals. (**E**) Frequencies of CD161^hi^ cluster relative to all T cell subsets. Females were plotted in red and males in blue. Red dash box delineated the healthy females. (**F**) Top enriched pathways of CD161^hi^ cluster in Reactome Pathway Database ranked by false discovery rate (FDR, - log10 scale). Data were plotted as mean ± standard error. Significance was determined by Kruskal-Wallis test (E).

### Sex-Specific Differences of Circulating MAIT cells in COVID-19

To analyze our scRNA-seq dataset for potential sex differences in circulating CD161^hi^ cells, we first sought to examine for phenotypic heterogeneity within this population. To do this, we performed a focused sub-cluster analysis, which generated 3 distinct clusters (**Fig. 3A**). However, the added resolution revealed a cluster that expressed *TRDC*, encoding the constant region of the δ chain expressed by γδ T cells (**Fig. 3B**) and thereby excluded from subsequent analyses. By contrast, the other two clusters had higher *KLRB1* expression, as well as *TRAV1-2* (**Fig. 3B**), therefore referred to here as MAITα and MAITβ clusters. Of note, these 2 clusters make up approximately 80% of CD161^hi^ PBMCs, which is consistent with the previous report of circulating MAIT cell frequencies (*25*). Our results showed that the MAITα cluster possessed upregulated genes associated with cytotoxic T cells (*GNLY, CD8A, CD8B*), migration/adhesion (*CXCR4, ITGB2*), and cytokine signaling (*IRF1*, *B2M*, *NFKBIA*, *JUNB, FOS*) (**Fig. 3C** and **table S4**). The MAITβ cluster was enriched for genes of ribosomal proteins, apoptosis (*BAX*, *STUB1*) and the linker histone H1 associated with apoptosis (*HIST1H1C, HIST1H1D, HIST1H1E*) (**Fig. 3C** and **table S4**). Gene enrichment analysis further supported a functional dichotomy for α and β clusters. Whereas MAITα was enriched with several immune process pathways (e.g., IFN-γ, and IL-4 and IL-13 signaling, as well as antigen processing and presenting), MAITβ was enriched in cellular responses to external stimuli, metabolism of RNA, viral infection, and programmed cell death, but not immune processes (**Fig. 3, D** and **E**, and **table S5**). Hence our results suggest MAIT cell heterogeneity, with the MAITα signature representing an immunologically poised/active phenotype, while the MAITβ signature represents a stressed/apoptotic phenotype.

**Fig. 3.**
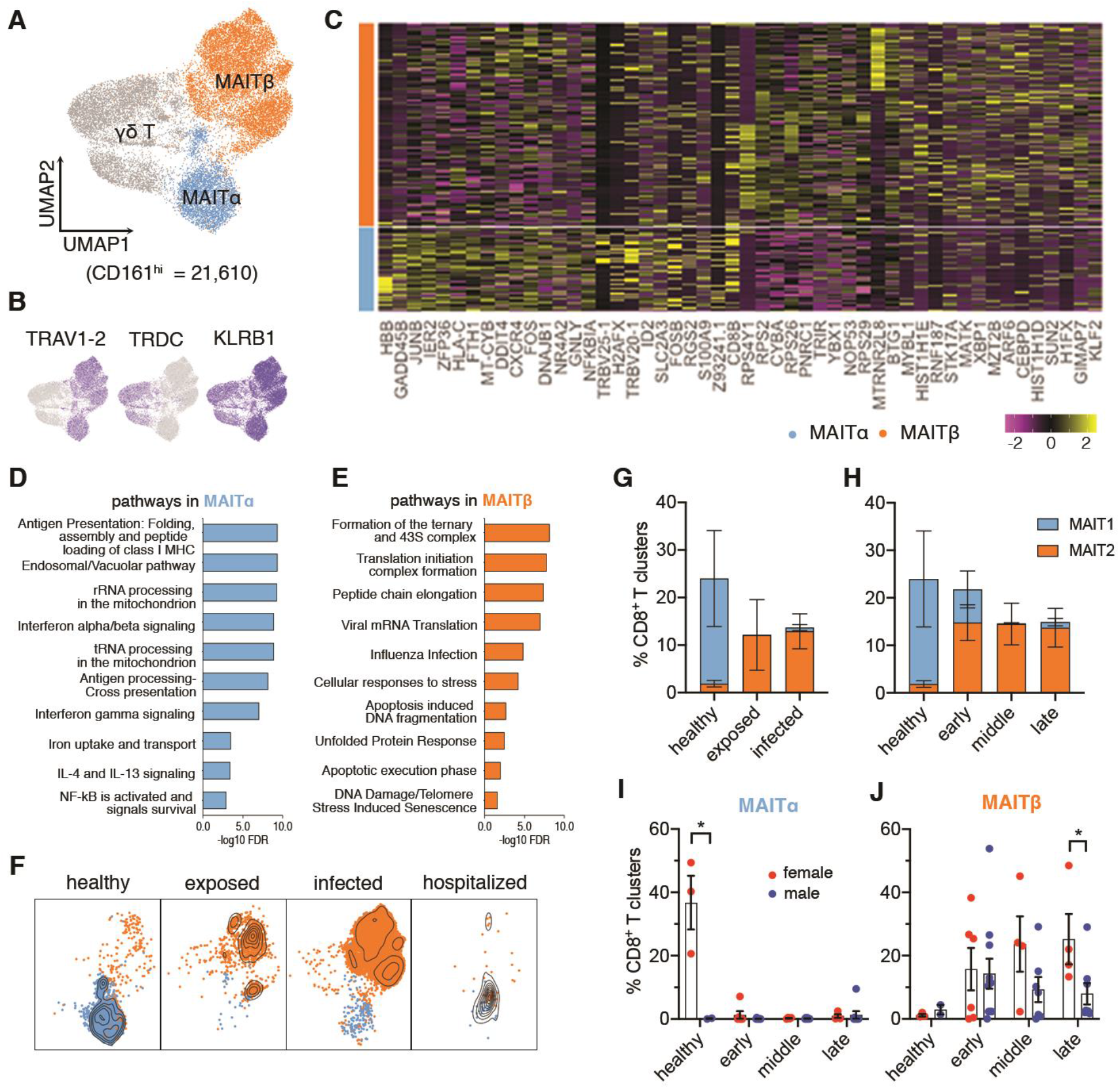
Heterogeneity and Dynamics of Circulating MAIT cells across Sex in COVID-19. (**A**) Sub-clustering of CD161^hi^ cells (n = 21,610) showing two MAIT clusters and one γδ T cluster. (**B**) Marker gene expression of three CD161^hi^ clusters. (**C**) Heatmap of top 25 discriminative genes between MAITα and MAITβ clusters. Expression level was scaled by Z-score distribution. (**D** and **E**) Representative top enriched pathways of MAITα and MAITβ in Reactome Pathway Database (ranked by false discovery rate, - log10 scale). Top 100 DEGs ranked by fold change between MAITα and MAITβ were used for this analysis. (**F** and **G**) UMAP visualization of MAIT cluster changes (F) and their frequencies (G) with disease severity. (**H**) Frequencies of MAIT clusters grouped by time post symptom onset. (**I** and **J**) Sex differences of MAIT clusters as shown in H. Data were plotted as mean ± standard error (G-J). Significance was determined by Mann Whitney test (I). *p<0.05.

Last for this series of experiments, we sought to determine the dynamics of the two phenotypically distinct clusters by sex over the COVID-19 disease course. By first grouping our data by severity, we found that MAITα was the major phenotype in healthy individuals, while MAITβ predominated in exposed and infected groups (**Fig. 3, F** and **G**). There was a noted exception for hospitalized patients (**Fig. 3F**), bearing very few cells as seen in our flow cytometry data, consistent with lymphopenia that occurs in severe COVID-19 (*24*). We then grouped our data by time post symptom onset, as we detailed earlier with our flow cytometry data. Results showed that relative to healthy subjects, MAITα percentages were lower in early, middle, and late timepoints, whereas MAITβ demonstrated the converse (**Fig. 3H).** When stratified by sex, we found that MAIT cell frequencies were higher in healthy females (**Fig. 3, I** and **J**), corroborating our flow cytometry results. These cells in healthy females were skewed toward the MAITα cluster, whereas the few cells present in healthy males consisted mostly of MAITβ (**Fig. 3I**). However, this difference was lost in exposed/infected setting, where both sexes were comprised mostly of MAITβ (**Fig. 3J**). Nonetheless, MAITβ percentages were statistically greater in females in late disease (**Fig. 3J**), which reflects the increased MAIT cells during late infection in females as shown in our flow cytometry findings. Regarding expression of *CD69*, a T cell activation marker, we did not observe major differences across cluster or sex, but did observe elevated expression in the hospitalized group (**Fig. S6, A** to **D**). This possibly suggests an altered MAIT cell response in hospitalized patients (*25–27, 35*). In short, these results reveal sex specific MAIT cell differences at the quantitative and phenotypic levels in health and COVID-19.

### Respiratory Tract MAIT Cell Responses Differ by Sex in COVID-19

To assess potential sex-specific differences in MAIT cells at the tissue level in COVID-19 patient airways, we utilized published scRNAseq datasets of bronchoalveolar lavage fluid (BALF) (*36*) and of nasopharyngeal swab (NPS) (*37*). Beginning our analysis with the BALF dataset, we identified the MAIT cell cluster by expression of *TRAV1-2*, *CD3D*, *KLRB1* and *SLC4A10* (**Fig. 4, A** to **C**). With this annotation, we found a significant increase (p=0.0188) of MAIT cells in COVID-19 patients relative to normal controls and a higher MAIT cell frequency (p=0.0332) in females relative to males among COVID-19 subjects (**Fig. 4D**). This detection of increased MAIT cells in female BALF, along with the drop in these cells we observed in female peripheral blood, suggest a female dominant extravasation of MAITs in COVID-19.

**Fig. 4.**
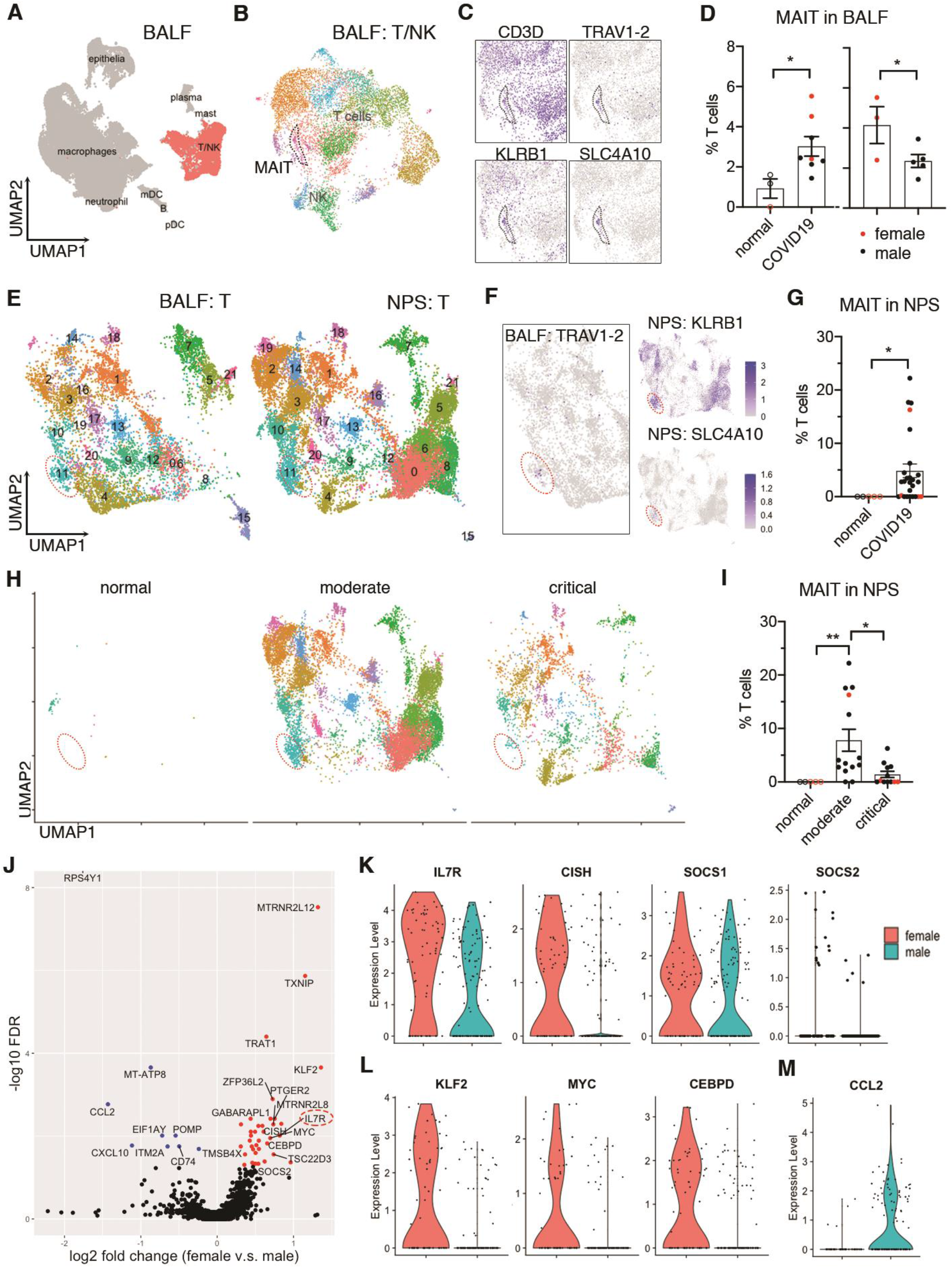
MAIT Cell Differences by Sex in Airway Tissue Samples of COVID-19 Patients. (**A** and **B**) Clustering analysis of scRNA-seq data from COVID-19 BALF dataset with subtracted T and NK cells (*36*). (**C**) MAIT cell cluster indicated by marker genes. (**D**) Frequencies of MAIT cells in BALF between normal and COVID-19 subjects (left) and across sex within COVID-19 subjects (right). (**E**) Integrated clustering analysis of NPS with BALF using Seurat 3. (**F**) Referenced MAIT cluster in NPS by the expression of TRAV1-2 in BALF and indicated marker genes in NPS. (**G**) Frequencies of MAIT cells in NPS from healthy and COVID-19 subjects. (**H** and **I**) Visualization (H) and frequencies (I) of MAIT cells in NPS grouped by disease severity. (**J**) Volcano plot showing of DEGs of BALF MAIT cells between sex with fold change and FDR. (**K** to **M**) Expression of DEGs in IL-7 signaling (**K**), transcriptional factors (**L**) and CCL2 (**M**). Data were plotted as mean ± standard error (**D, G, I**) with females in red and males in black. Significance was determined by unpaired a one-tailed student’s t-test (D), Kruskal-Wallis test with Dunn’s post hoc test (I) and Mann Whitney test (G): *p<0.05, **p<0.01.

To perform the same analysis with NPS samples, we integrated the T cell data from NPS with those from BALF to identify MAIT cells in NPS (**Fig. 4, E** and **F**), since TCR genes were not aligned in the NPS dataset (*37*). With this annotation, we quantified MAIT cell frequencies in the NPS dataset, again observing a significant increase (p=0.0139) in COVID-19 patients (**Fig. 4G**). We also analyzed the data across severity, observing a significant increase (p=0.0038) in moderate subjects relative to the normal and a decrease (p=0.0366) relative to critical COVID-19 subjects (**Fig. 4 H, I**). We could not perform the same analysis by sex due to insufficient number of female samples (**Fig. 4 I**). Nonetheless, these results match the reduced circulating MAIT cell frequencies seen in our hospitalized subjects, together suggesting that a lymphopenic state which occurs in severe COVID-19 impacts MAIT cells, consistent with other reports (*25–27, 38*).

In a final experiment, we sought to characterize MAIT cell transcriptomes by sex in the BALF and NPS datasets and determine whether these cells resembled α and β phenotypes we identified in circulating MAIT cells. Cluster analysis was not warranted here given low cell numbers in these datasets. Instead, we leveraged gene modules derived from our respective α and β clusters of circulating MAIT cells. We found that MAIT cells in BALF and NPS data were largely skewed toward the β module, with minimal sex differences (**Fig. S7A**). However, when we directly examined differentially expressed genes (the DEGs between sex, we were able to detect sex differences associated with α and β phenotypes. Specifically, in BALF, we found increased *IL7R* expression in females (**Fig. 4J**) and other IL-7 signaling associated genes (*CISH* and *SOCS1*) (**Fig. 4K**). Given the critical role of this signaling in T cell survival, we explored additional pathway genes, finding that female MAIT cells had upregulated anti-apoptotic genes (*BCL2* and *FOXP1*) and downregulated pro-apoptotic genes (*BAX* and *CASP3*) (**Fig. S7, B** and **C**). Also observed in female cells was upregulated anti-proliferative genes (*CDKN1B* and *BTG2*) (**Fig. S7D**). These patterns matched MAITα gene changes in our PBMC dataset. We were able to find other sex differences, including increased expression of several transcription factors (*KLF2*, *MYC*, and *CEBPD*) (**Fig. 4L**). Conversely, male cells had higher expression of *CCL2* (**Fig. 4 M**), which has been linked to COVID-19 immunopathology (*39*). In short, our results infer sex differences at the qualitative level in COVID-19, with female MAIT cells possessing a pro-survival and immunologically active phenotype.

## DISCUSSION

Despite the knowledge of sex differences in the immune response as an underlying factor in COVID-19 disease outcomes, the sexual dimorphic responses of MAIT cells, an unconventional T cell population deemed important in this disease, remained unknown. We now demonstrate that MAIT cells in females are quantitatively and qualitatively more robust in the SARS-CoV-2 setting, potentially helping understand the immunological reasons for reduced COVID-19 susceptibility in females.

Our finding that MAIT recruitment to airway tissues may be more robust in COVID-19 females was aided first by our observation of higher circulating MAIT cell frequencies in females in the healthy setting. This difference can be explained by the rate of physiological aging-related attrition of MAIT cells that is substantially less pronounced in female blood (*40–42*). The resultant higher frequencies in circulation enabled us to readily uncover the precipitous percentage drop we saw with MAIT cells relative to exposed/infected females. In trying to elucidate the potential cause of this drop, we considered two possible scenarios: 1) lymphopenia and 2) extravasation, which are not necessarily mutually exclusive. For the former, it is accepted that lymphopenia is associated with severe COVID-19 infections (*24, 32, 43, 44*), which which agrees with our observations in our hospitalized group (comprised of 77.8% intensive care patients). Similarly, lymphopenia could partially explain the reduction in MAIT cells described by Jouan *et al* in a study of male-dominated samples from critically ill COVID-19 patients (*27*), though extravasation also likely occurred. In our study, however, we demonstrated that circulating MAIT frequencies drop in our infected outpatient group. As these subjects were not critically ill, our findings point to extravasation as a major reason for the sex-specific drop in circulating MAIT frequencies. The same pattern may also exist in several other studies (*25, 26, 35*). For example, while Parrot *et al* (*25*) also demonstrated that circulating MAIT cells are reduced in moderate COVID-19 patients relative to healthy subjects in aggregated data, it is possible that the healthy female frequencies contributed to reaching the statistical difference. Further supporting our conclusion, we were able to show with publicly available scRNA-seq data from COVID-19 BALF samples (*36*) that females in that study had an increased MAIT cell percentage relative to males, allowing us to conclude that MAIT cell extravasation during COVID-19 may be quantitatively more robust in females.

Our results also suggest that MAIT cells may be qualitatively superior in females, with respect to anti-viral immune activity in COVID-19. Leading us to this conclusion, our scRNA-seq analysis of patient PBMCs revealed two distinct clusters of MAIT cells, referred to here as MAITα and MAITβ. The α cluster was enriched for various immune pathways, such as IFN-γ signaling, inferring a capacity for anti-viral immune function. In contrast, the β cluster was enriched for cell stress and apoptosis pathways, inferring a frail phenotype roughly similar to a previously described population of double negative MAIT cells (*45, 46*). We showed in the healthy setting that MAIT cells in females were skewed toward the α cluster, whereas males comprised the β cluster. Though from these results it could be presumed that the α cluster should be overrepresented in COVID-19 airways of females, this was not the case in the BALF. However, we reasoned that such a finding would be very difficult to make for two main reasons. First, extravasated MAIT cells with an α-phenotype would be restricted to the early wave of recruitment, since circulating cells are almost completely skewed to the β module in exposed/infected individuals. Second, a certain level of transcriptional reprogramming would occur upon immune cell extravasation into the tissue and potentially again upon accessing the alveolar space. Still, we were able to show in BALF that certain gene patterns remained consistent with the α signature in females versus males. In addition, our finding that female BALF samples had quantitatively more MAIT cells gives further credence that differences revealed in blood would likewise extend to the tissue.

In summary, we conclude that MAIT cells in females are quantitatively and qualitatively distinct from males and we surmise that this distinction provides a protective advantage in the SARS-CoV-2 setting. Indeed, females in general tend to have elevated frequencies of circulating MAIT cells, also gleaned by large independent studies with European (*40*), South Korean (*41*) and Chinese populations (*42*). Further supporting this argument, it has now been recorded that adult COVID-19 fatality rates trend less in females at all ages across 39 different countries, including in North America, Europe, and Asia where MAIT cell frequencies trend higher in females (*2*). These points also argue against the possibility that an immunologically more robust MAIT cell response has a net negative effect, for example, by immunological misfiring (*20*) or cytokine storm related immunopathology (*39*). However, one open question that our findings now raise is whether males in our study, which had greater circulating CD8+ memory T cells, would instead have an advantage in the reinfection setting or following vaccination.

Future studies are needed to explore this question, and to better understand sex differences in MAIT cells both in general and in COVID-19.

## MATERIALS AND METHODS

### Ethics statement

This study and relevant protocols were approved by the Institutional Review Boards of Duke University Health System (DUHS) ?. All procedures were performed in accordance with the Declaration of Helsinki, applicable regulations, and local policies.

### Participants in this study

In-patients (hospitalized) and out-patients (infected) with confirmed infection of SARS-CoV-2 were identified through the DUHS and enrolled into the Molecular and Epidemiological Study of Suspected Infection (MESSI, Pro00100241). The RT-PCR testing for SARS-CoV-2 was performed at either the North Carolina State Laboratory of Public Health or at clinical laboratories of the DUHS. The exposed group, who closely contacted with COVID-19 patients, presented negative PCR test and negative serology test during longitudinally sampling from the first visit to at least 2 months after, typically 0, 7, 14, and 28 days relative to enrollment. Initial severity scores of individuals were recorded through a self-reporting survey on 38 defined symptoms related to COVID-19 plus “other” when enrolled. The exposed group (average score = 9.71) showed a lower severity scores compared with infected (out-patients) group (average score = 18.16). The hospitalized patients presented severe disease symptoms with breath difficulty, cough, fever or chest pain when enrolled, and 77.8% of them for this study required intensive care unit (ICU) care. All COVID-19 patients were also longitudinally sampled with serology test from enrollment to convalescent phase. Healthy donors were enrolled in 2019 (Duke IRB Protocol Pro00009459) with no diagnosis or symptoms consistent with COVID-19 or other respiratory illness. Written informed consent was obtained from all subjects or legally authorized representatives. Patient Demographics are summarized in Table 1.

### Collection of peripheral blood mononuclear cells (PBMCs)

PBMC cells were prepared using Ficoll-Hypaque density gradient method. Briefly, peripheral whole blood was collected in EDTA vacutainer tubes and processed within 8 hours. Blood was diluted 1:2 in PBS then layered onto the Ficoll-Hypaque in 50 ml conical tube and centrifuged at 420g for 25 min.

Buffy coat was collected and washed with D-PBS by centrifugation at 400g for 10min. Cell pellets were resuspended in D-PBS and washed again. PBMCs were assessed for viability and cell count using Vi-Cell automated cell counter (Beckman-Coulter). PBMCs were adjusted to 10×10^6^ cells/ml in cryopreservation media (90% FBS, 10% DMSO) and aliquoted into cryopreservation vials on ice. Cells underwent controlled freezing at −80℃ using CoolCell LX (BioCision) for 12-24 hours, then transferred to liquid nitrogen vapor phase.

### Sample processing for flow cytometry and single cell RNA-sequencing (scRNA-seq)

Counts and cell viability of thawed PBMCs were measured by Countess II after a wash with DMEM 10% FBS. The cell viability of hospitalized patients ranged from 70-80% whereas all other samples exceeded 80% viability. An additional dead cell removal step (Miltenyi Biotec) was conducted on hospitalized PBMC samples prior to aliquot for scRNA-seq. To perform scRNA-seq, 200,000 cells per sample were aliquoted, spun down, resuspended in 30 μl PBS supplemented with 0.04% BSA and 0.2U/ μl RNase inhibitor and counted using Countess II.

### Panel and Staining for Flow Cytometry

Approximately 0.5-2 × 10^6^ cells per cryopreserved sample were stained for flow cytometry analysis. Antibody titrations used in this study were previously established by Cytek Biosciences with slight modifications (see Table S2 for flow panel information). All staining procedures were performed at room temperature. PBMCs were stained with live/dead Blue (Thermofisher) for 15 min, washed with FACS-EDTA buffer and spun down at 1500 rpm for 5min. Samples were resuspended with Brilliant Stain Buffer Plus (BD Biosciences) and sequentially stained with anti-CCR7 for 10 min, the chemokine receptor mix for 5 min, anti-TCR gamma/delta for 10 min and the surface receptor mix for 30 min. After incubation, PBMCs were washed with FACS-EDTA buffer and spun down at 1500 rpm for 5min. Samples were fixed with 1% PFA in PBS for 20 min, spun down and resuspended in FACS-EDTA buffer.

### 36-color Full Spectrum Flow Cytometry

Samples were acquired using a four-laser Cytek Aurora Spectral Flow Cytometry System. Single color controls for spectral unmixing were done with PBMCs from healthy control blood and UltraComp eBeads (ThermoFisher). Raw data were unmixed and further analyzed using either FlowJo for manual gating or Omiq (https://www.omiq.ai) for clustering visualization and analysis.

### High-dimensional data analysis of flow cytometry data

Uniform Manifold Approximation and Projection (UMAP) and FlowSOM clustering analyses were performed on Omiq (https://www.omiq.ai), using equal random sampling of 3000 live CD45+ singlets. from each FCS file. The UMAP plot was generated with the parameters of 15 neighbors and 0.4 minimum distance. All markers in flow panel were used for analysis except live/dead and CD45.

### ScRNA-seq using 10x Genomics platform

10x Genomics Single Cell 5’ v1 chemistry was used to generate Gel Bead-In Emulsions (GEM), and perform post GEM-RT cleanup, cDNA amplification, as well as library construction. An agilent DNA ScreenTape assay was used for quality control. Libraries were pooled and sequenced to saturation or 20,000 unique reads per cell on average using an Illumina NovaSeq6000 with 150-bp paired-end reads.

### Processing and quality control of scRNA-seq

Raw sequencing data were initially processed with 10x Genomics Cell Ranger pipelines (V3.1.0). Briefly, BCL files were demultiplexed to generate FASTQ files. FASTQ files were aligned with STAR aligner to the human genome reference GRCh38 from Ensemble database. Feature barcode processing and UMI counting were then performed according to the standard workflow. (QC summary after sequencing). The following criteria were applied as quality control of single cells from all individual samples. Cells that had fewer than 1000 UMI counts or 500 genes, as well as cell that had greater than 10% of mitochondrial genes were removed from further analysis. Genes that were expressed by fewer than 10 cells were also excluded. After filtering, a total of 424,080 cells with 18,765 gene features were kept for the downstream analysis.

### Dimensionality reduction and clustering analysis

The filtered gene-barcode matrix was analyzed using Seurat 3 (*28*). All the procedures were conducted with the default parameters unless otherwise specified. Briefly, data were first normalized using log transformation and adjusted with a scale factor of 10,000. The top 2,000 variable genes were identified, and percentages of mitochondrial genes were regressed out when scaling data. Principle component analysis (PCA) was performed using these top variable genes, and top 25 principle components (PCs) were selected for graph-based clustering with Shared Nearest Neighbor (SNN) and visualization in UMAP. The resolution was set to 0.35 to identify major immune cell subsets in PBMCs. Sub-clustering of CD161hi T cells (21,610 cells) was also performed using the analytic pipeline mentioned above with two modifications: top 10 PCs were used, and the resolution was set to 0.1 to identify MAIT cell clusters.

### Differential gene expression analysis

Differentially expressed genes (DEGs) were identified using Seurat 3 (FindAllMarkers or FindMarkers Functions) with either ‘wilcox’ for all cluster markers or ‘DESeq2’ (*47*). Randomly downsampled data with 100,000 cells were used to find all markers of PBMC clusters. A gene was considered significant with adjusted p-value or false discovery rate (FDR) < 0.05. DEGs results of all PBMCs and MAIT cells are listed in Table S3 and S4.

### Pathway enrichment analysis

Top 100 DEGs of MAIT clusters were used for pathway enrichment analysis using Reactome Pathway Database (https://reactome.org). A pathway was considered significantly over-presented with FDR < 0.05. The full pathway enrichment results are summarized in Table S5.

### Inference of ligand-receptor interactions between T cells and monocytes

Ligand-receptor interactions between T cells and monocytes were inferred using CellPhoneDB (*33*). PBMC scRNA-seq data were randomly downsampled to 50,000 cells and T and monocyte clusters were extracted based on the expression of their lineage markers. CellPhoneDB was with default parameters (https://github.com/Teichlab/cellphonedb). The inferred interactions are considered significant when p-value < 0.05.

### Integration of BALF and NPS dataset

Publicly available scRNA-seq data of BALF (*36*) and of NPS (*37*) were downloaded and processed using Seurat 3 as previously described (*28*). All T cell clusters, were extracted from both dataset and integrated via Single Cell Transform (SCT) method in Seurat 3. Top 3,000 variable features were selected for the integration. Dimensionality reduction was conducted using PCA and UMAP embedding of the top 100 PCs. Clusters were visualized at a resolution of 0.8 after constructing a SNN graph using the first 50 PCs.

### Calculations of the feature scores in MAIT cells

The DEGs between MAIT1 and MAIT2 were used to generate their feature scores as previously described (*48*). The feature scores were calculated using AddModuleScore function in Seurat 3. MAIT cells from different single cell dataset were plotted with MAIT1 feature and MAIT2 feature for visualization.

### Statistical analysis

Data normality and homogeneity of variance were assessed using Kolmogorov-Smirnov test and Bartlett’s test, respectively. Due to the distribution and variance of human data, non-parametric statistical tests were favorably used throughout this study unless otherwise specified. Mann Whitney U test was used for two-group comparisons, and Kruskal-Wallis with post hoc Dunn’s test was used for comparisons of three groups and more. Spearman’s correlation efficiency was used to quantify the correlation of the ranked disease severity (from healthy as 1, to hospitalized as 4). To adjust p-values for multiple hypothesis testing, FDR correction was performed using the Benjamini-Hochberg procedure when appropriate. Two-tailed tests were used unless otherwise specified. A p-value or FDR < 0.05 is consider statistically significant. Graphical data of quantifications presented throughout are expressed as the means ± SEMs and were plotted using Graphpad Prism 8. Other graphs in this study were generated using either the corresponding analytic packages or R package ggplot2.

### Data availability

All clinical metadata of participants and samples in this study are included in Table S1. Data will be shared upon the acceptance of this manuscript. Publicly available scRNA-seq data of BALF (*36*) were downloaded from GEO with the accession number GSE145926, and the count data of NPS (*37*) were downloaded from https://doi.org/10.6084/m9.figshare.12436517. All of the raw fcs files and all scripts used for data analysis are available to share per request.

## Acknowledgements

We would like to thank Monica DeLay and Patrick Duncker (Cytek Biosciences) for their help with spectral flow cytometry, and Chris Ciccolella and Geoff Kraker (Omiq, Inc). We would also like to thank Maria Miggs, Deborah Murray, Tyffany Locklear, Robert Rolfe, Jack Anderson, Allison Fullenkamp, Raul Louzuo, Thad Gurley and Julie Steinbrink for their work, as well as the support from Durham Veterans Affairs Health Care System and Duke Regional Hospital.

## Supplementary Materials

**Table S1. Clinical Metadata of Participants and Samples in this study**

**Table S2. List of Panel Reagents for 36-color Spectral Flow Cytometry**

**Table S3. List of DEGs of PBMC Clusters and MAIT cells**

**Table S4. List of DEGs of MAIT Subsets**

**Table S5. List of Enriched Reactome Pathways of MAIT Subsets**

**Figure S1.**
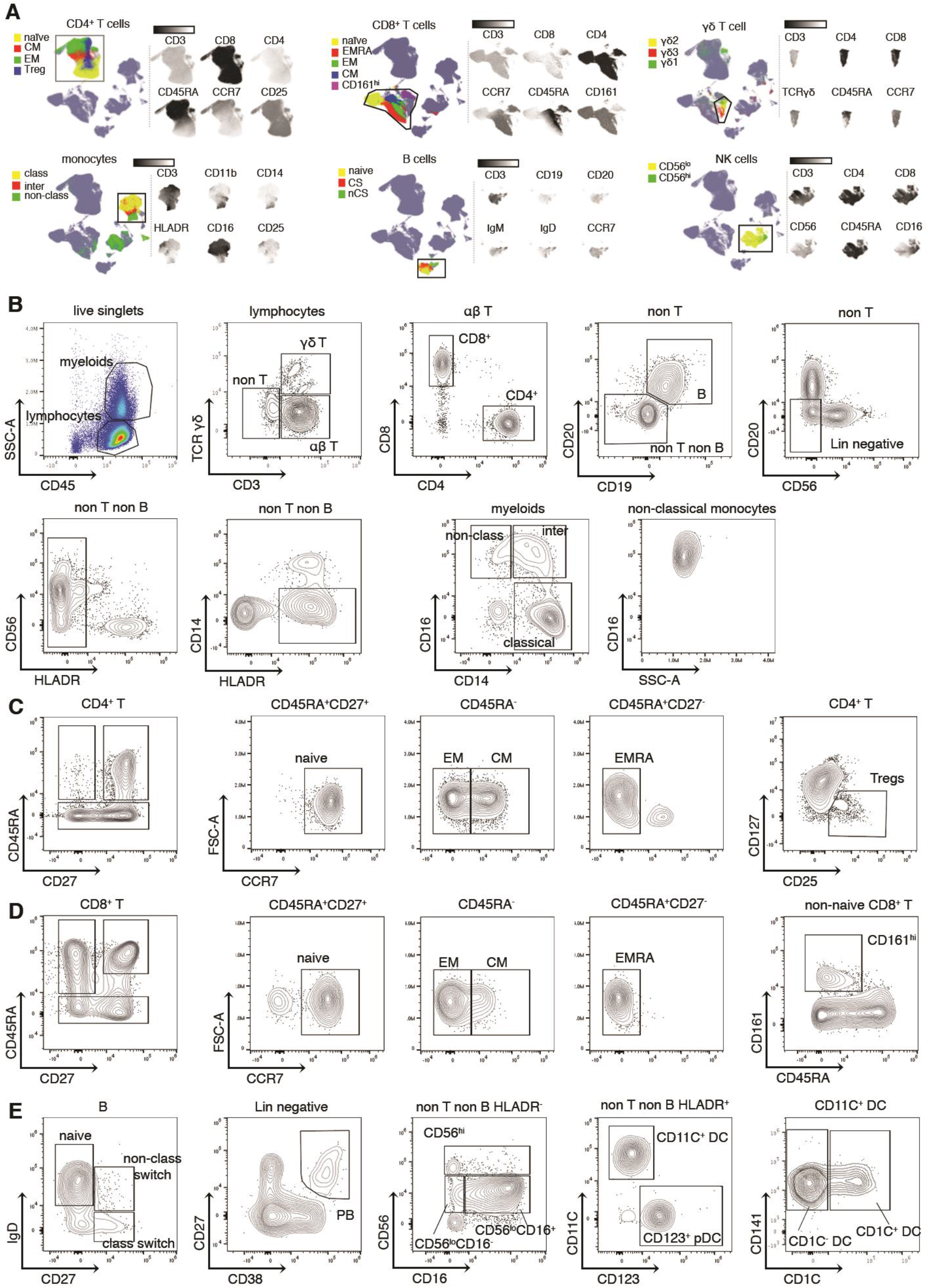
Annotation of Immune Subsets and Gating Strategy of PBMCs from COVID-19 patients Using 36-color Flow Cytometry. (**A**) Annotation of subsets for CD4 T, CD8 T cells, γδ T cells, monocytes, B cells and NK cells as shown in fig. 1B. Left-hand side in each panel is the UMAP with the indicated immune cell population (box) and the respective subsets annotated by color (legend). Right-hand side is feature plots of the indicated population showing relative expression of each marker (black= negligible; white = positive). (**B**) Gating strategy of immune cell subsets in PBMC samples.

**Figure S2.**
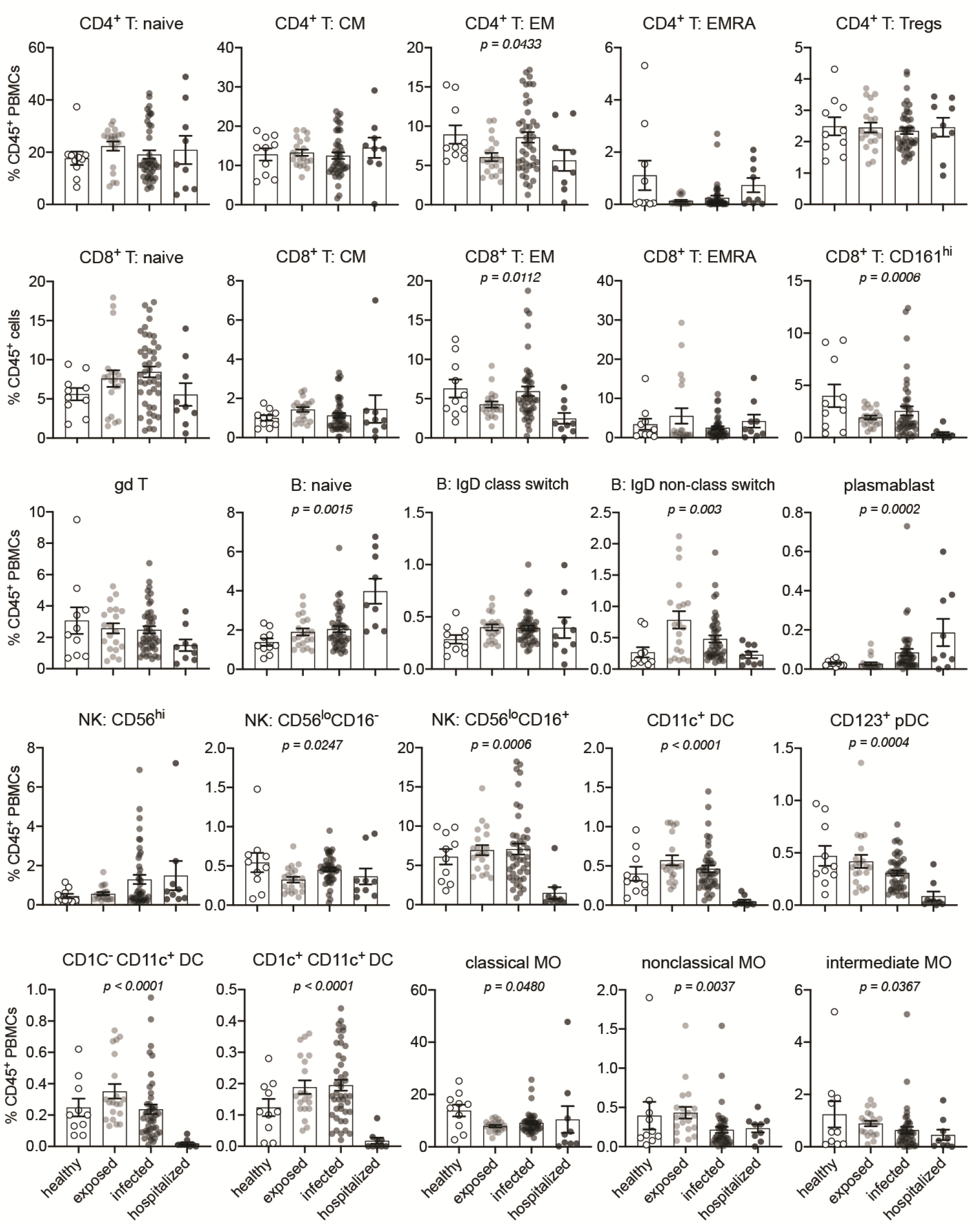
Frequency Profiling of Immune Subsets in COVID-19 PBMCs among Different Disease Severity groups. Gating strategy was shown in Fig. S1B. All data were plotted as percentage of CD45^+^ PBMCs. Significance was determined by Kruskal-Wallis test.

**Figure S3.**
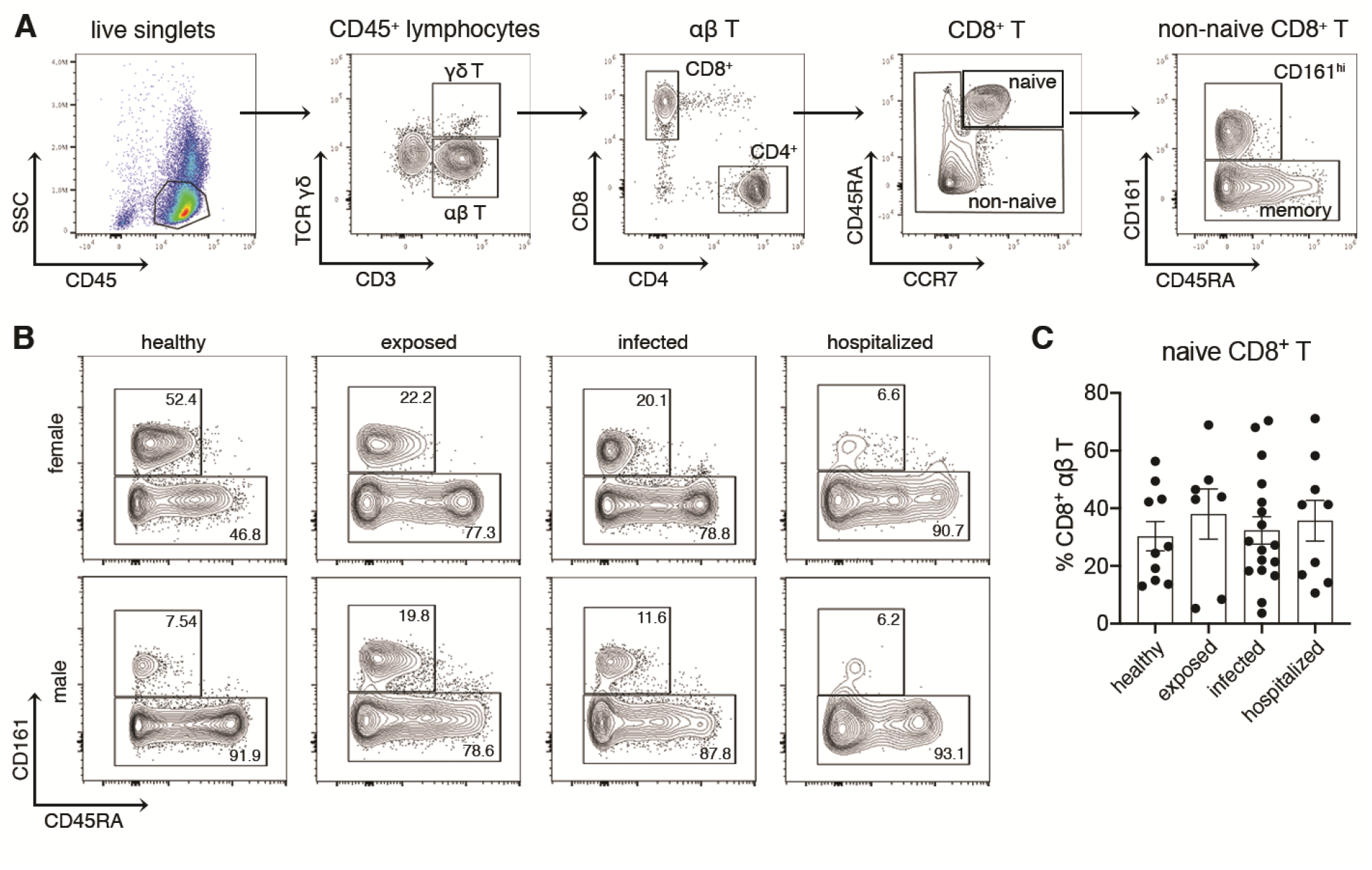
Sex-Specific Changes of CD8^+^ CD161^hi^ and Memory T Cells in COVID-19 PBMCs by Flow Cytometry. (**A**) Gating strategy of CD8+ CD161hi, memory and naive T cells (αβ). (**B**) Representative flow plots of CD8^+^ CD161^hi^ and memory T cells from female and male individuals with disease severity. (**C**) Frequencies of naïve CD8 T cells in different severity groups as indicated. Significance was calculated by Kruskal-Wallis.

**Figure S4.**
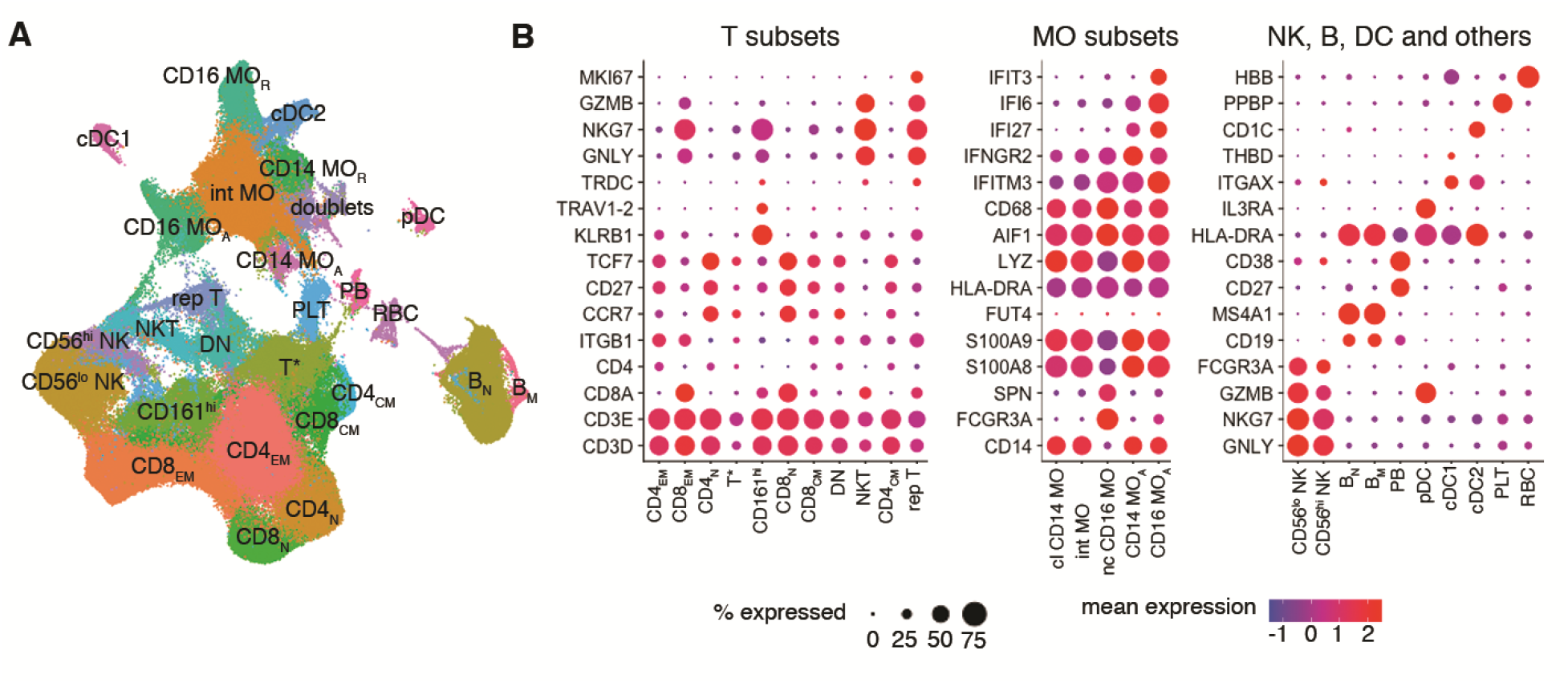
Characterization of Immune Subsets in COVID-19 PBMCs by scRNA-seq. (**A**) High resolution clustering of PBMCs shown in Fig.2A. (**B**) Expression of marker genes used to annotate individual clusters. cl, classical; int, intermediate; nc, non-classical; CD14 MOA, activated CD14 monocytes; CD16 MOA, activated CD16 monocytes; PB, plasmablasts; PLT, platelets.

**Figure S5.**
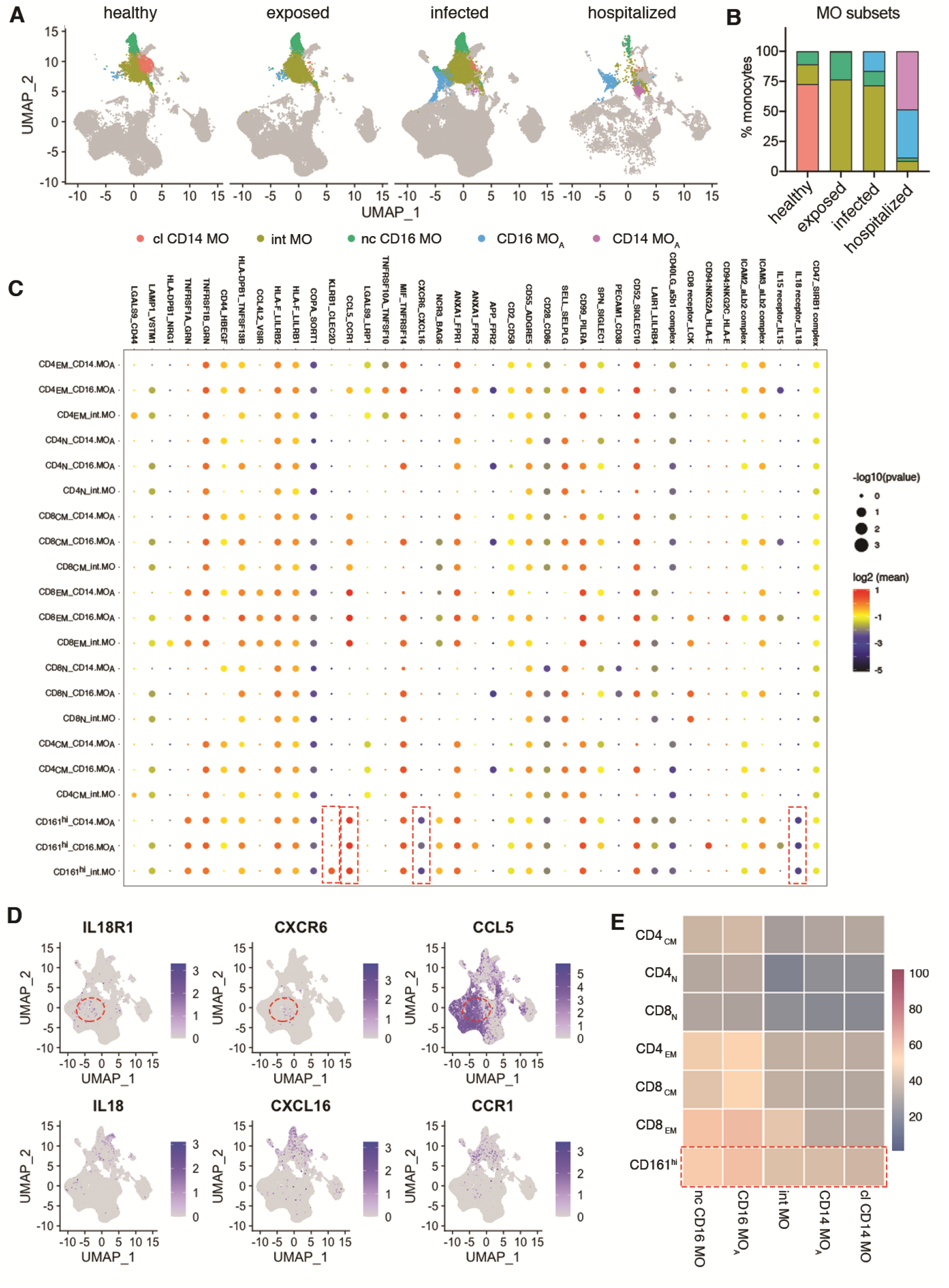
Interaction Inference of Receptors and Ligands between Monocytes and Major T Cell Subsets. (**A** and **B**) Visualization (**A**) and percentage distribution (**B**) of different monocyte subsets identified in Fig.S4A. Three monocyte subsets represent resting classical CD14 (cl), non-classical CD16 (nc), and intermediate (int) monocytes as seen in healthy subjects. Two monocyte subsets are associated with interferon signaling and COVID-19 patients with differential expression of CD14 and CD16, referred to as activated CD14 and activated CD16 monocytes (CD14 MOA and CD16 MOA, respectively). (**C**) Overview of selected ligand–receptor interactions inferenced by CellPhoneDB in COVID PBMC single cell dataset. Red dash box delineated the specific interaction of CD161^hi^ T cells with monocytes. P values and scales are indicated by circle size and colors, respectively. (**D**) Expression of representative ligand and receptor pairs between MAIT and monocytes as indicated. (**E**) Heatmap of interaction counts between major T cell and monocyte subsets.

**Figure S6.**
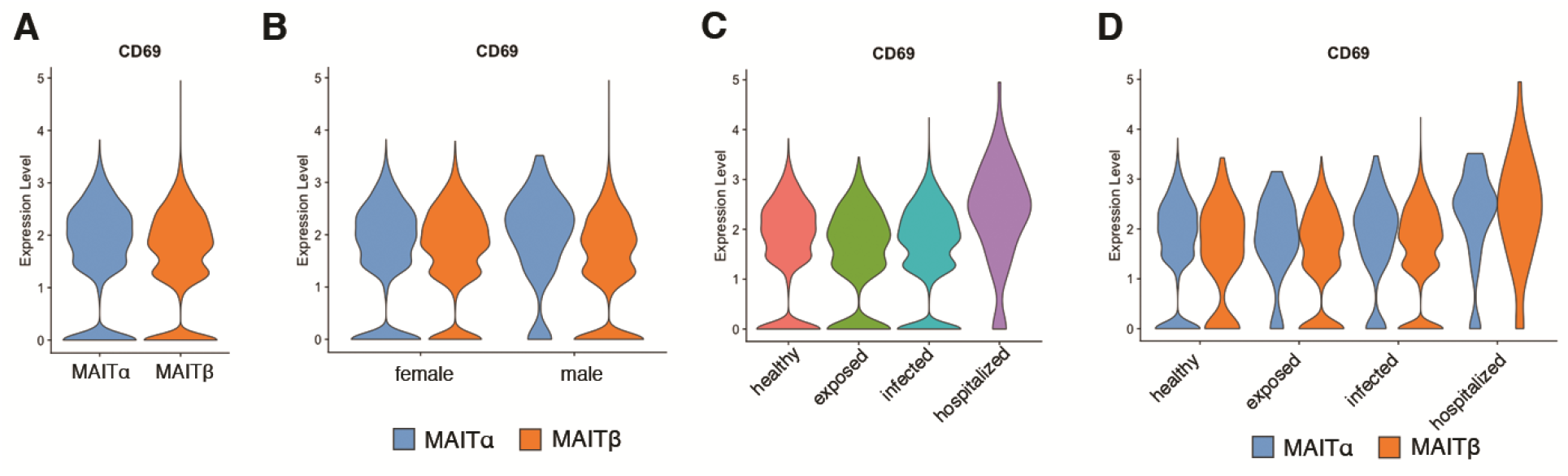
Expression of CD69 by circulating MAIT Clusters in COVID-19. (**A** to **D**) Data are grouped by MAIT clusters (A), sex (B), disease severity (C) as well as clusters and severity (D).

**Figure S7.**
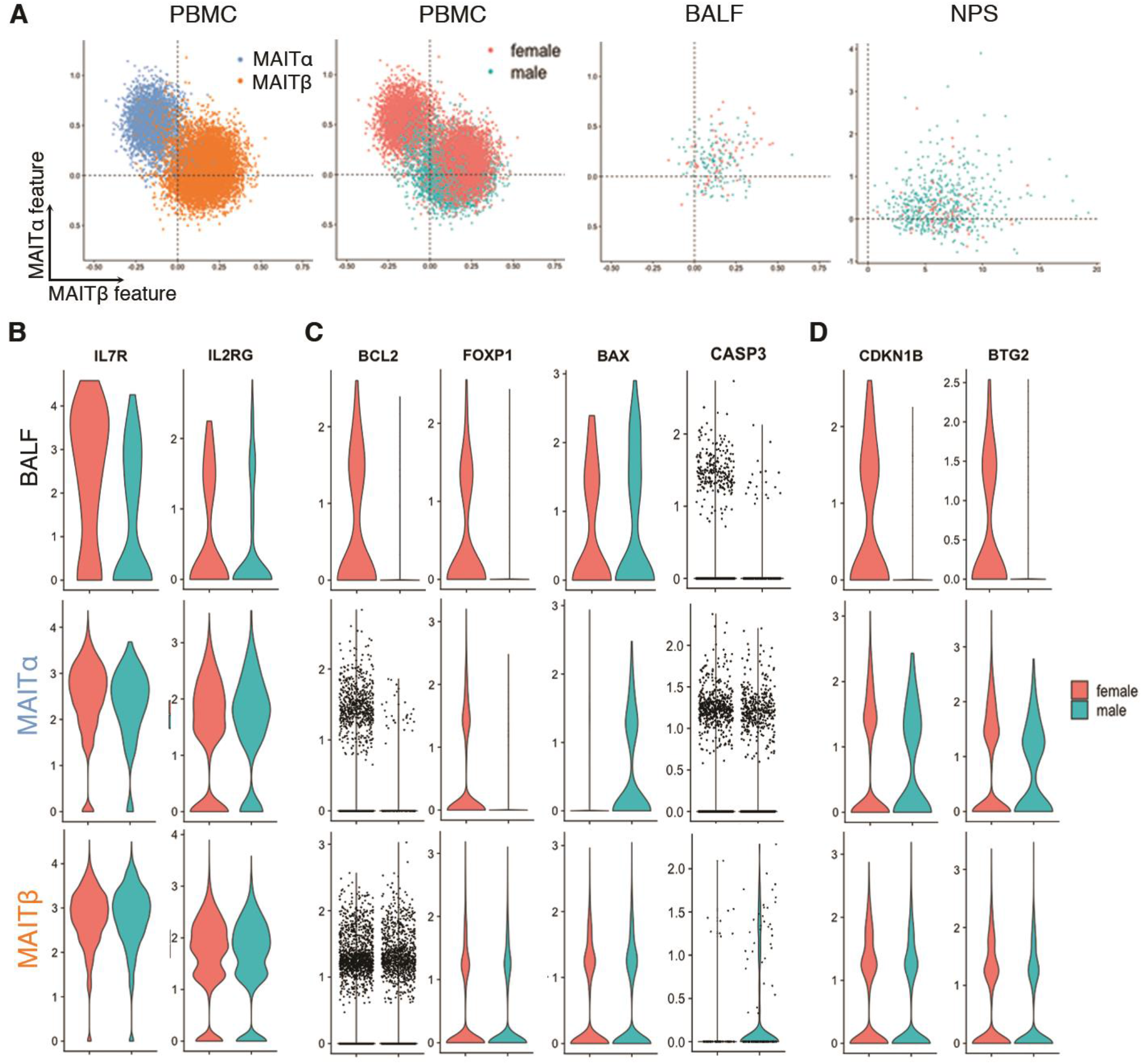
Comparisons of MAITα and MAITβ Features in MAIT Cells from peripheral blood and airway tissue samples. (**A**) Estimation of MAITα and MAITβ features of individual cells from PBMC and BALF dataset based on the expression of MAITα gene (y-axis) and MAITβ gene sets (x-axis). Differentially expressed genes between MAITα and MAITβ were used as two modular features, respectively. (**B to D**) Expression of IL7 receptor and its co-receptor genes (B), apoptosis-related genes (C), proliferation-related genes by MAIT cells from PBMC and BALF dataset (D).

